# Mitigation of Parkinson’s Disease Pathology in *C. elegans* by Marine Bacterium *Kocuria rhizophila* via Ferroptosis Suppression

**DOI:** 10.64898/2026.08.25.746916

**Authors:** Simran Singh, Anusree Damodaran, Neeraj Kumar, Pushplata Yadav, Mukesh Pasupuleti, Sonia Verma

## Abstract

Parkinson’s disease (PD) is a progressive neurodegenerative condition characterized by the loss of dopaminergic (DA) neurons and alpha-synuclein aggregation, with ferroptosis playing a critical pathological role. This study investigated the neuroprotective potential of *Kocuria rhizophila* strain CDMP12, a marine bacterium isolated from the Gulf of Mannar, India, using *Caenorhabditis elegans* models of PD. Dietary supplementation with *K. rhizophila* (CDMP12) significantly preserved DA neuron structure, rescued neuro-sensory and motor deficits, and attenuated both alpha-synuclein expression in the *C. elegans* models. Transcriptomic and qRT-PCR analyses revealed that CDMP12 systematically suppressed ferroptosis by significantly downregulating iron and lipid regulatory genes such as *smf-3*, *ftn-1*, and *acs-4*, while upregulating the protective antioxidant gene *gpx-1*. Furthermore, BODIPY staining demonstrated that CDMP12 treatment markedly reduced lipid peroxidation, lowering the oxidized-to-non-oxidized lipid ratio in PD worms. Collectively, these findings identify *K. rhizophila* (CDMP12) as a promising marine-derived neuroprotective candidate that mitigates PD-associated pathology, accompanied by reduced alpha-synuclein burden, preservation of DA neuronal function, and attenuation of ferroptosis-associated molecular and lipid peroxidation signatures.

**Highlights:** 1. Marine bacterium *K. rhizophila* preserves dopaminergic neurons in *C. elegans* Parkinson’s disease models.

2. *K. rhizophila* restores dopamine-dependent behaviors in UA44 worms.

3. *K. rhizophila* reduces alpha-synuclein expression.

4. *K. rhizophila* induces widespread transcriptomic remodelling

5. *K. rhizophila* normalise the expression of key ferroptotic genes and reduces lipid peroxidation; implicating ferroptosis as a central target.

## 1. Introduction

Parkinson’s disease (PD) represents one of the most rapidly escalating neurological conditions globally, with estimates predicting that its prevalence will exceed 12 million cases by 2040 ^1,2^. Pathologically, PD is characterized by the degeneration of dopaminergic (DA) neurons within the substantia nigra pars compacta (SNpc), culminating in classic motor impairments, such as resting tremors, as well as non-motor symptoms, including cognitive decline ^3^. Although existing clinical interventions like dopamine agonists and monoamine oxidase-B inhibitors offer meaningful symptomatic relief, they do not alter the underlying trajectory of the disease and frequently induce debilitating side effects ^4,5^. Therefore, shifting the therapeutic paradigm away from simple symptom mitigation toward targeting the core mechanisms of neurodegeneration is essential for developing true disease-modifying treatments.

A promising candidate in this space is ferroptosis, an iron-dependent form of programmed cell death driven by toxic lipid peroxidation ^6,7^. Pathological features characteristic of PD, including iron overload, elevated lipid peroxides, and marked glutathione depletion, strongly implicate ferroptotic signaling in DA neuronal loss ^8–12^. Crucially, targeting this cascade has yielded encouraging pre-clinical results: the small-molecule ferroptosis inhibitor ferrostatin-1 rescues DA neurons and alleviates motor deficits in numerous in vitro and in vivo PD models ^13–16^. Taken together, these data highlight ferroptosis modulation as a viable and highly promising avenue for neuroprotective therapy in PD.

The marine environment has emerged as an under-explored bio-resource for discovering structurally diverse neuroprotective agents. Over the past two decades, more than 50 natural compounds isolated from marine bacteria, fungi, and macroalgae have demonstrated promising therapeutic activity across preclinical and clinical evaluation ^17,18^. For example, NP7, a secondary metabolite derived from *Streptomyces* sp. crosses the blood-brain barrier, mitigates oxidative stress-induced neurotoxicity, and dampens microglial activation ^19,20^. Likewise, piloquinones A and B from *Streptomyces* sp. CNQ-027 act as potent inhibitors of monoamine oxidase-B ^19,21^. Notably, marine-derived natural products are also advancing into clinical trials: docosahexaenoic acid has been shown to alleviate depressive symptoms in PD patients (NCT01563913), inosine was evaluated for its capacity to elevate neuroprotective urate levels (NCT02642393), and ganglioside GM1 demonstrated early clinical promise (NCT00037830) ^22^. Collective evidence underscores the marine biome as a rich, yet largely unexplored, reservoir for next-generation PD neuroprotective interventions.

Model organisms remain fundamental to anti-Parkinsonian drug discovery due to their conserved biological pathways and shared disease mechanisms with humans. Among these systems, *Caenorhabditis elegans* serves as an exceptionally tractable model for studying aging and neurodegeneration, owing to its short lifespan and genetic homology with human ^23,24^. Transgenic *C. elegans* strains expressing human alpha-synuclein reproduce hallmark PD phenotypes, including age-dependent protein aggregation, selective DA neurodegeneration, and quantifiable DA-mediated behavioral deficits. Crucially, these nematodes have facilitated the discovery of key genetic modifiers and small molecules capable of protecting DA neurons ^23–25^. Consequently, *C. elegans* provides an efficient, high-throughput platform to screen marine bacterial therapeutics and elucidate translational neuroprotective targets for human application.

In this study, we investigated the neuroprotective potential of *Kocuria rhizophila* strain CDMP12, a marine bacterium isolated from seawater samples collected in the Gulf of Mannar, India. During an initial phenotypic screen of marine bacterial isolates, *K. rhizophila* (CDMP12) conferred protection against DA neurotoxicity in *C. elegans* models of PD, prompting its selection for detailed mechanistic investigation. While *K. rhizophila,* a Gram-positive actinobacterium has previously been characterized for its agricultural and environmental applications, such as enhancing plant growth under heavy metal and saline stress, as well as biosorbing heavy metals like cadmium and chromium from wastewater, its biomedical and therapeutic potential remains largely unexplored ^26–31^. We demonstrate that feeding on CDMP12 effectively rescues DA neurodegeneration, attenuates alpha-synuclein expression, and restores both motor and sensory behavioral functions in *C. elegans* PD models. These neuroprotective effects were accompanied by suppression of ferroptosis-associated molecular signatures, including reduced lipid peroxidation and altered expression of key iron- and lipid-regulatory genes. Collectively, our findings establish *K. rhizophila* (CDMP12) as a novel marine-derived candidate for neuroprotective therapy while further reinforcing ferroptosis inhibition as a target for PD intervention (**Grpahical Abstract**)

## 2. Materials and Methods

### 2.1. Isolating *Kocuria rhizophila* (CDMP12)

The isolation and molecular identification of *K. rhizophila* (CDMP12) (NCBI GenBank: KM677905.1) followed procedures established in our previous studies ^25,32^. In brief, seawater samples from the Gulf of Mannar were collected using specific transport media, serially diluted, and cultured on Zobell’s marine agar at 37°C for two days. Genomic DNA extraction (Sigma-Aldrich GenElute Kit, NA2110) and 16S rRNA gene PCR amplification using primers 63F and 1387R (**Table 1**) were carried out as previously described ^27^. Amplicons were sequenced via an ABI 3730XL system, and readouts were checked for quality and contig consistency using DNA Baser and DECIPHER prior to BLAST verification ^25,32–34^. The partial sequence of the 16S rRNA gene results obtained from the Standard Nucleotide Blast analysis were submitted to the NCBI accession number (NCBI GenBank: KM677905.1).

**Table 1:**
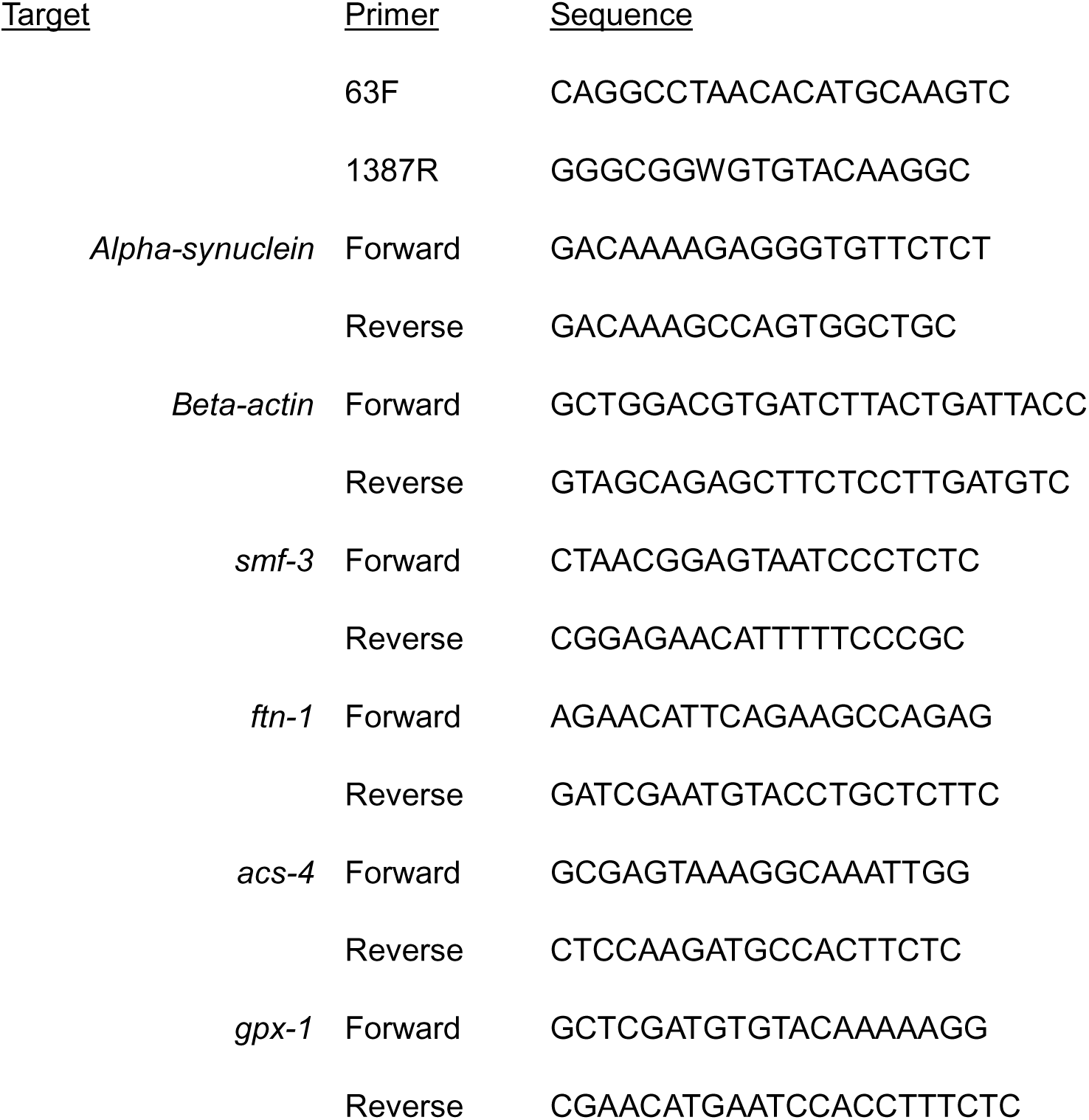
Details of Primers used in the study:

### 2.2. Maintaining *C*. *elegans* strains

We utilized three well-established *C. elegans* strains for this investigation: UA44 [baInl1 (P_dat-1_:: alpha-synuclein, P_dat-1_::GFP)]: Co-expresses human alpha-synuclein and GFP specifically in DA neurons, serving as a model for DA neurodegeneration and behavioral deficits.

BY250 [vtIs7 (P_dat-1_::GFP)]: Expresses GFP in DA neurons to function as a healthy control for UA44.

NL5901 [P_unc-54_:: alpha-synuclein::YFP + unc-119(+)]: Expresses human alpha-synuclein-YFP fusion protein within body-wall muscle cells to monitor protein aggregation.

The UA44 line was generously provided by Dr. Anoopkumar Thekkuveettil from the Sree Chitra Tirunal Institute for Medical Sciences and Technology (Thiruvananthapuram, India). All other strains were acquired from the *Caenorhabditis* Genetics Center (University of Minnesota, USA). Worms were cultured on standard nematode growth medium (NGM) plates seeded with *Escherichia coli* OP50 at 20°C ^25,35^. To generate age-synchronized population, gravid adults received standard hypochlorite treatment (bleaching) to harvest eggs. The resulting embryos hatched in M9 buffer, arresting development at the L1 larval stage over 16-18 hour incubation at 20°C ^25,35^. These synchronized L1 larvae were subsequently transferred onto experimental NGM plates seeded with either OP50 or CDMP12.

### 2.3. Preparing bacteria-seeded NGM plates

Cultivation of OP50 and CDMP12 was performed as previously described, using Luria–Bertani broth (GLR Innovations, Cat. No. GLRCM0054) and Zobell’s Marine Broth (HiMedia, Cat. No. M385-500G), respectively ^25^. Following overnight incubation at 37°C, cultures were pelleted at 6000 rpm, washed in M9 buffer, and resuspended to 2 mg/mL. NGM plates (60 mm) were seeded with 0.5 mL of the bacterial suspension and dried for 16 hours before adding L1 larvae. Progeny production was blocked by adding 0.1 mg/mL of fluoro-2′-deoxy-β-uridine (TCI, Cat. No. D2235) at the L4 stage. All behavioral, phenotypic, and molecular evaluations were performed on day three of adulthood, maintaining a consistent feeding schedule throughout. This specific time point was chosen because robust alpha-synuclein-related pathologies and DA degeneration are observable by day three in these PD models ^25,36^.

### 2.4. Assessing DA neuronal health

DA neuron integrity was assessed on day three of adulthood using fluorescence microscopy and as previously described, ^25^. Briefly, worms were collected and washed from plates using M9 buffer, mounted on 2% agarose pads, and immobilized with 30 mM sodium azide (Sigma, Cat. No. S2002). Fluorescence was visualized at 20× magnification on a Leica DMi6000 microscope. Neuron fluorescence intensity was quantified using ImageJ (NIH, Bethesda, MD) by tracing the DA neurons in the head region. Data represent a minimum of 30 worms per condition pooled from at least three biological replicates.

### 2.5. Assessing alpha-synuclein expression

Alpha-synuclein expression was evaluated in three-day-old adults by examining fluorescence in strain NL5901 ^25^. Briefly, nematodes were collected, washed with M9 buffer, and immobilized on 2% agarose 22pads. Worms were immobilised using 30 mM sodium azide and fluorescence images of the worms were captured at 10× magnification using a Leica DMi6000 fluorescence microscope. Fluorescence intensity was subsequently quantified using ImageJ software. Data represent a minimum of 30 worms per condition pooled from at least three biological replicates.

### 2.6. Quantifying pharyngeal pumping rate

Pharyngeal pumping rates were evaluated on day three of adulthood following established protocols ^25^. Briefly, video recordings of at least 30 seconds were captured for individual worms using a dissection microscope (Weswox Optik SZM-102) equipped with a camera. Pharyngeal contractions were manually counted over to calculate the pumping rate. Data represent 30 worms per condition pooled from at least three biological replicates.

### 2.7. Assesing 1-Nonanol avoidance

Avoidance behavior was assessed on day three of adulthood via nonanol repulsion assay ^25^. Nematodes were harvested and washed with M9 buffer before transferring at least 30 worms to the middle of four-quadrant 60 mm unseeded NGM plates. 1 µL of 1-nonanol (TCI, Cat. No. N0292) was spotted onto two diagonally opposite quadrants. After incubating the plates for 45 minutes at 20°C, the worms were counted in each quadrant to compute the repulsion index.

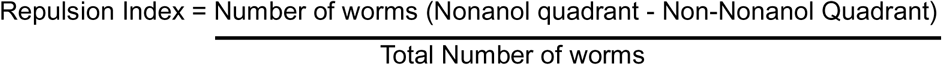

Each condition was tested across a minimum of three independent biological replicates.

### 2.8. Thrashing assay

Locomotor capacity was evaluated by quantifying head thrashing frequency following established protocols ^37^. To remove adherent bacteria, adult worms were gently transferred onto unseeded NGM plates and allowed to crawl freely. Individual animals were then placed into 60 mm empty plates filled with 1 mL of M9 buffer. After a 1-minute acclimation period to stabilize swimming behavior, head thrashes were observed under a stereomicroscope and recorded over a 30-second interval. Data represent 30 worms per condition pooled from at least three biological replicates.

### 2.9. Assessing Basal Slow Response

The basal slowing response was evaluated as previously described ^38^. Briefly, least 30 worms from each experimental group were transferred to seeded or unseeded NGM plates. Behavioral assessments were conducted using a stereomicroscope, where the number of body bends was counted over a 20-second interval to quantify crawling speed. The basal slowing response was calculated using the following formula:

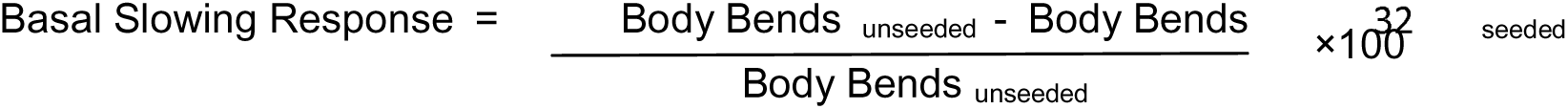

Data represent 30 worms per condition pooled from at least three biological replicates.

### 2.10. RNA isolation and quantitative real-time PCR (qRT-PCR)

On day three of adulthood, total RNA was extracted from harvested worms using Trizol reagent (Sigma-Aldrich, Cat. No. T9424) combined with freeze-thaw cycles and vortexing, followed by phenol:chloroform:isoamyl alcohol separation and isopropanol precipitation ^25^. Roughly 2 µg of RNA was reverse-transcribed into cDNA with the GoScript Reverse Transcription System (Promega, Cat. No. A5001). qRT-PCR was performed on a Bio-Rad CFX96 thermocycler using SYBR Premix Ex Taq (TaKaRa Bio Inc., Cat. No. RR420A). Gene expression levels were normalized to beta-actin and calculated via the 2^−ΔΔCt^ method. Primer sequences are provided in Table 1.

### 2.11. BODIPY staining

To measure lipid peroxidation on day three of adulthood, worms were stained with the oxidation-sensitive dye BODIPY 581/591 (Invitrogen, Cat. No. D3861) ^25^. A 5 mg/mL DMSO stock was diluted to 1 µg/mL in M9 buffer. Harvested worms were incubated in 500 µl of working solution for 90 minutes at room temperature with gentle agitation. Following washes, animals were mounted, and 10× fluorescence images were acquired using a Leica DMi6000 microscope. Because the dye shifts from red (reduced) to green (oxidized) upon peroxidation, oxidation levels were quantified as the ratio of green-to-red fluorescence intensity using ImageJ. Data represent a minimum of 10 worms per condition pooled from at least three biological replicates.

### 2.13. Transcriptomic analysis

Sample processing, RNA sequencing, and downstream analyses followed previously described methods ^25^. Briefly, worms (UA44 (OP50) and UA44(CDMP12)) were grown in six independent biological replicates (∼500 worms/replicate). On day three of adulthood, triplicates were pooled to create two biological replicates (∼1500 worms each) per condition. Pellets were washed, snap-frozen, and processed by Biokart Genomics Lab (Bengaluru, India). Total RNA was extracted with the RNeasy Mini Kit (Qiagen) (RIN > 7.0), and libraries prepared via the NEBNext Ultra II RNA Library Prep Kit were sequenced on an Illumina HiSeq platform. Sequencing data were analyzed on Galaxy: reads were quality-checked with FastQC, trimmed using Trimmomatic, aligned to the *C. elegans* ce11 genome via HISAT2, and quantified using featureCounts. Differential expression was evaluated using LIMMA-voom, setting thresholds at adjusted *p* ≤ 0.05 and |log_2_ fold-change (FC)| ≥ 0.5.

### 2.13. Functional enrichment analysis

To evaluate functional enrichment, gene lists were analyzed using Database for Annotation, Visualization, and Integrated Discovery (DAVID) platform (https://david.ncifcrf.gov/) for Gene Ontology (GO) terms and Reactome pathways, applying a significance threshold of p ≤ 0.05 ^39^. GO categories were grouped into biological processes, cellular components, and molecular functions. Data visualization was performed in RStudio (version 1.3.959; https://rstudio.com/) using ggplot2 to construct bubble plots representing the enriched categories ^40^.

### 2.14. Statistical analysis

Statistical analyses were carried out in GraphPad Prism 8.0 (GraphPad Software, La Jolla, CA, USA). Results are presented as mean ± standard deviation. Differences between groups were assessed via Student’s t-test, considering p ≤ 0.05 as statistically significant.

## 3. Results

### 3.1. *K. rhizophila* (CDMP12) protects DA neurons against degeneration in the UA44 strain

PD is characterized by the progressive loss of DA neurons within the SNpc ^41^. Using *C. elegans* PD model, UA44, we evaluated the effect of feeding CDMP12 on DA neuron integrity on day three of adulthood using fluorescence microscopy. The UA44 (OP50) worms exhibited a decrease in DA neuron fluorescence (11.96 ± 0.58) compared to the healthy control, BY250 (OP50) worms (19.26 ± 0.85; p < 0.0001), validating the neurodegenerative phenotype. Feeding UA44 worms on CDMP12 significantly preserved DA neuron fluorescence (16.67 ± 0.74; p < 0.0001) relative to UA44 (OP50). This restoration indicates that *K. rhizophila* (CDMP12) exerts a protective effect against DA neurodegeneration and mitigates PD pathology (**Figure 1A and B**).

**Figure 1.**
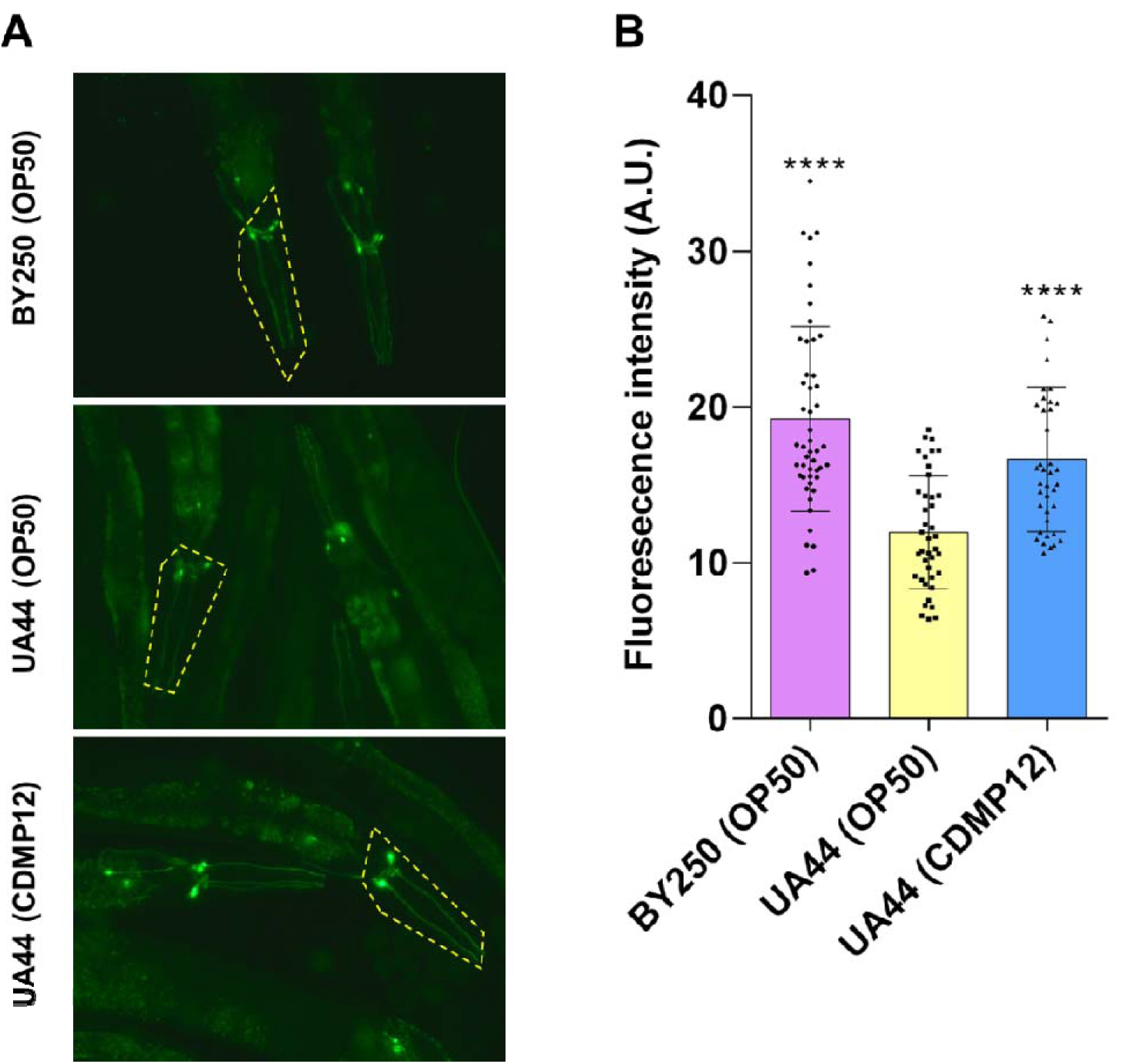
*K. rhizophila* (CDMP12) preserves DA neurons in the *C. elegans* PD model, UA44. (A) Head-region fluorescence micrographs from day t adults of BY250 (OP50), UA44 (OP50), and UA44 (CDMP12) worms (20X magnification). Yellow dashed outlines delineate the quantified head re encompassing both DA neuronal soma and processes. Intact DA structures in BY250 (OP50) contrast sharply with the severe degeneration seen in U (OP50) and CDMP12 feeding preserves neuronal integrity. (B) Quantification of DA fluorescence intensity via ImageJ. Relative to BY250 (OP50), U (OP50) worms exhibit lower fluorescence intensity. Supplementation with CDMP12 significantly restores DA fluorescence intensity in UA44 (n ≥ 30 worm condition). Data are presented as mean ± SD; ****p < 0.0001.

### 3.2. *K. rhizophila* (CDMP12) prevents neuro-sensory and motor deficits in UA44

Following structural neuron preservation, we assessed whether *K. rhizophila* (CDMP12) restores dopamine-dependent behaviors in UA44 strain. Using a 1-nonanol repellent assay to indirectly evaluate functional DA signaling, we observed a severe avoidance deficit in UA44 (OP50) worms (-0.07 ± 0.03) relative to BY250 (OP50) (-0.56 ± 0.02; p < 0.001) ^25,42,43^. CDMP12 feeding rescued this aversive behavior in UA44 (-0.50 ± 0.07; p < 0.01), indicating functional maintenance of DA-driven chemosensation (**Figure 2A**).

**Figure 2.**
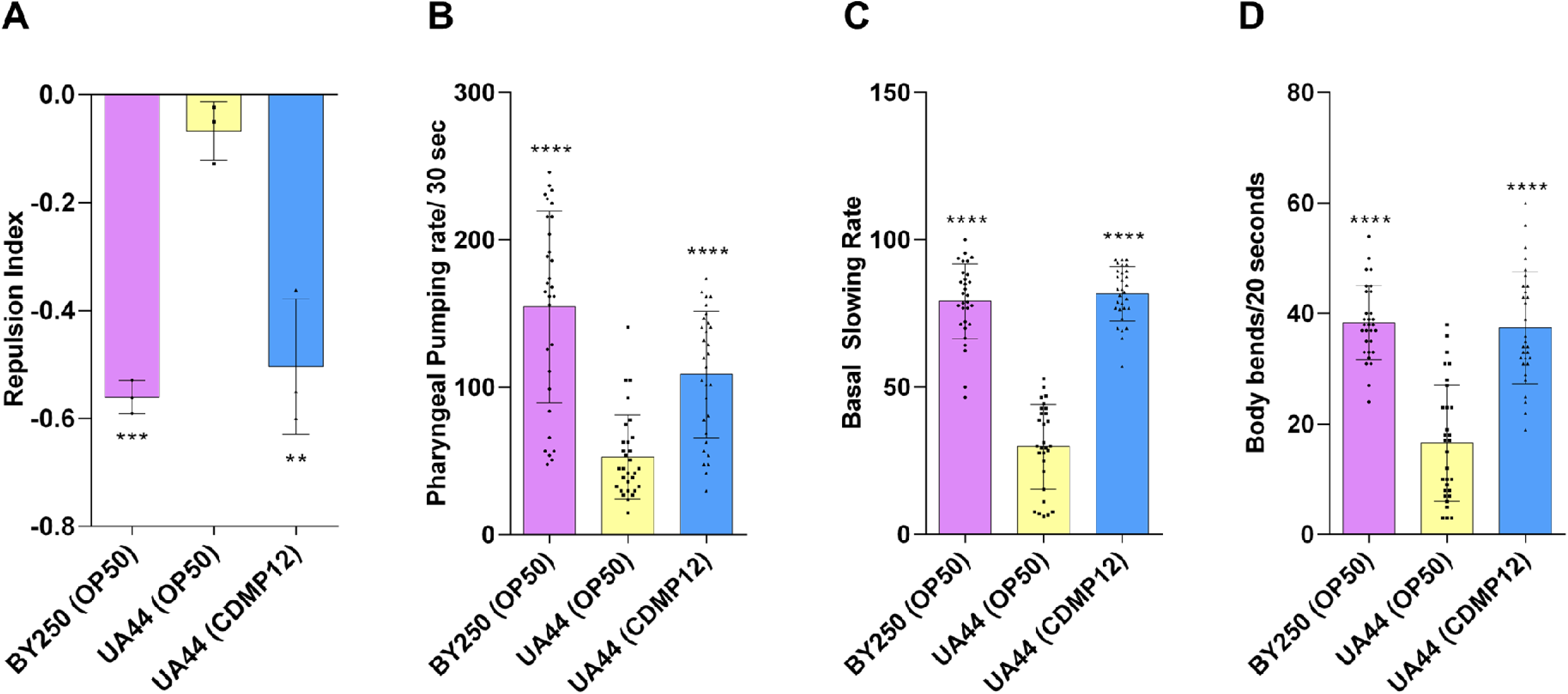
***K. rhizophila* (CDMP12) prevents dopamine-dependent neuro-sensory and motor deficits in the *C. elegans* PD model, UA44.** Quantification of repulsion index in the 1-nonanol avoidance assay. UA44 (OP50) exhibited a reduced aversive response compared to BY250 (O indicating impaired dopamine-dependent chemotaxis. CDMP12 supplementation significantly improved the repulsion index in UA44 worms, confirmation functional restoration of dopamine-driven behavior (n = 3 independent biological replicates; ≥30 worms per condition per replicate). (B) Quantification pharyngeal pumping rate (pharyngeal contractions per 30 seconds). UA44 (OP50) worms showed a decline in pumping frequency relative to BY250 (O reflecting alpha-synuclein–associated neuromuscular toxicity. CDMP12 diet significantly rescued pharyngeal pumping rate in UA44 worms (n ≥ 30 worm condition). (C) Quantification of the basal slowing response. UA44 (OP50) worms displayed a significantly attenuated slowing rate compared to BY250 (OP50), indicating compromised dopamine signaling. Dietary supplementation with CDMP12 significantly restored the basal slowing response in UA44 worms to control levels animals (n ≥ 30 worms per condition). (D) Assessment of locomotory capacity via body bend frequency in liquid (body bends per 20 seconds). UA44 (OP50) worms showed a severe reduction in body bend rate relative to BY250 (OP50), reflecting alpha-synuclein–induced motor impairment. CDMP12 supplementation markedly rescued locomotory performance in UA44 animals (n ≥ 30 worms per condition). Data are presented as mean ± SD; **p<0.01***p < 0.001, ****p < 0.0001.

We next evaluated neuromuscular toxicity by monitoring pharyngeal pumping ^25,44,45^. UA44 (OP50) worms displayed a decline in pumping rate/ 30 seconds (53.00 ± 5.19) relative to BY250 (OP50) (154.8 ± 11.88; p < 0.0001). This deficit was markedly attenuated in UA44 (CDMP12) (109.0 ± 7.84; p < 0.0001), demonstrating resistance to neuromuscular toxicity (**Figure 2B**).

To further confirm DA circuit protection, we tested the food-sensing BSR ^38^. UA44 (OP50) animals exhibited a blunted slowing response (29.88 ± 2.67) relative to BY250 (OP50) (79.26 ± 2.329; p < 0.0001), which was restored in UA44 (CDMP12) (81.70 ± 1.67; p < 0.0001), highlighting preserved neurochemical signaling (**Figure 2C**).

Lastly, liquid thrashing assay was used to assess locomotory performance ^46^. UA44 (OP50) worms displayed reduced number of body bends/ 20 seconds (16.60 ± 1.93) compared to BY250 (OP50) (38.37 ± 1.23; p < 0.0001). CDMP12 supplementation rescued thrashing frequency in UA44 (37.47 ± 1.85; p < 0.0001), establishing comprehensive behavioral protection (**Figure 2D**).

### 3.3. *K. rhizophila* (CDMP12) suppresses alpha-synuclein expression in NL5901 strain

Pathological accumulation and aggregation of alpha-synuclein represent a key driver of neurodegeneration in PD ^47^. To evaluate whether *K. rhizophila* (CDMP12) alters this pathological hallmark, we monitored alpha-synuclein accumulation in NL5901 transgenic worms. NL5901 (OP50) exhibited robust alpha-synuclein expression, characterized by intense fluorescence (90.38 ± 4.08), reflective of an aggregation-prone phenotype. In contrast, CDMP12 supplementation significantly attenuated fluorescence intensity (66.31 ± 2.90; p < 0.0001), demonstrating a reduction in protein accumulation (**Figure 3A and B**).

**Figure 3.**
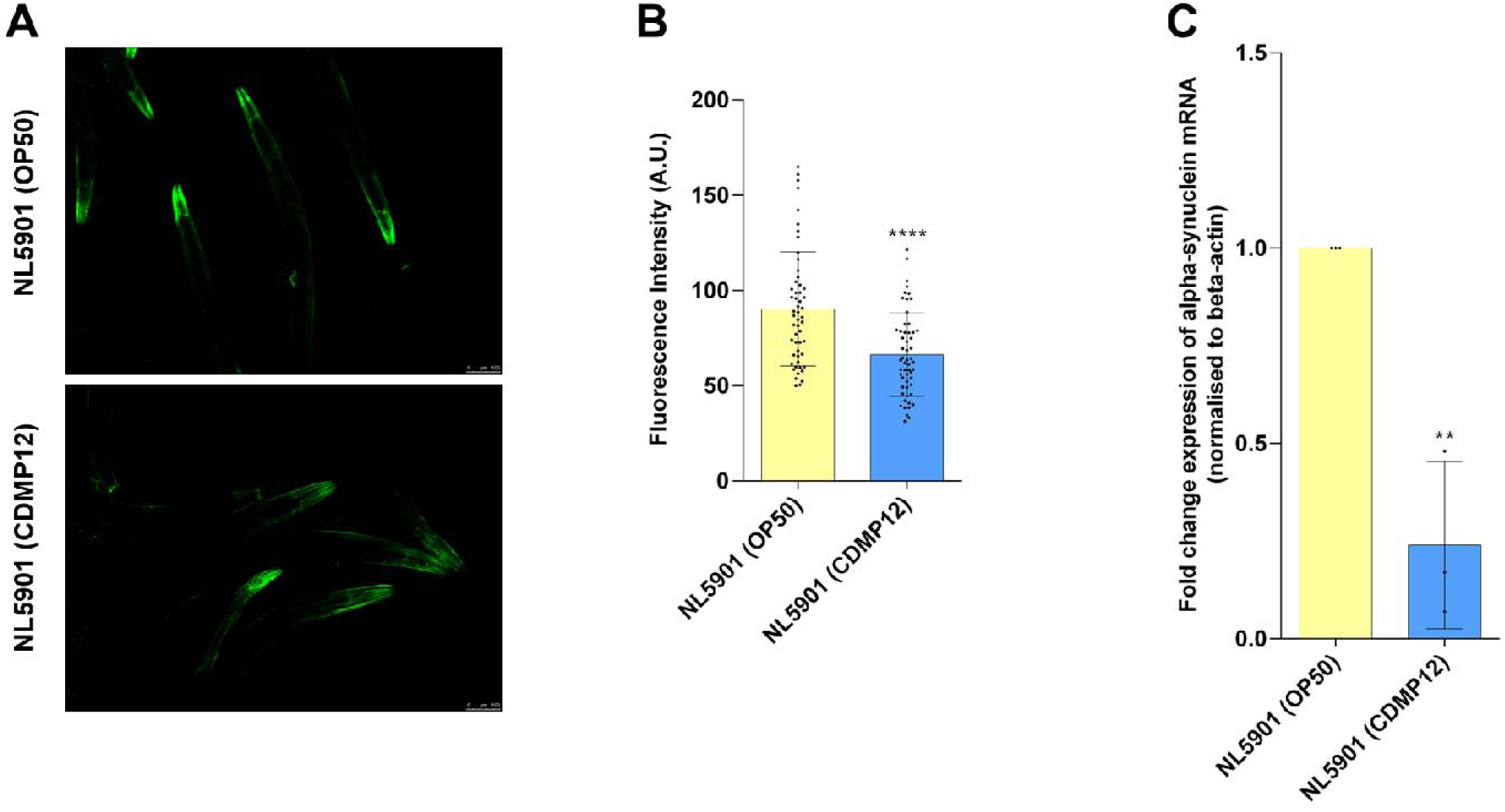
*K. rhizophila* (CDMP12) attenuates alpha-synuclein expression in the *C. elegans* PD model, NL5901. (A) Representative fluorescence micrographs of NL5901 worms expressing human alpha-synuclein::YFP in body-wall muscle. NL5901 (OP50) worms exhibit pronounced fluoroscence, w is reduced following CDMP12 supplementation (10X magnification). (B) ImageJ quantification of alpha-synuclein::YFP fluorescence intensity. NL5901 (O worms displayed enhanced fluorescence, whereas NL5901 (CDMP12) worms showed a significant reduction in fluorescence intensity, indicating suppressing of protein accumulation (n ≥ 30 worms per condition) (C) qRT-PCR quantification of alpha-synuclein mRNA transcript levels, normalized to beta-actin. NL (CDMP12) worms exhibited a marked downregulation of alpha-synuclein transcripts relative to OP50-fed controls (n = 3 independent biological replication >500 worms per condition per replicate). Data are presented as mean ± SD; **p < 0.01, ****p < 0.0001.

To determine whether this protein-level decrease stems from altered gene expression, we quantified alpha-synuclein mRNA transcript abundance. NL5901 (CDMP12) worms displayed a marked downregulation of alpha-synuclein transcripts (0.24 ± 0.12; p < 0.01) relative to NL5901 (OP50) (**Figure 3C**). Taken together, these data indicate that *K. rhizophila* (CDMP12) exerts its protective effects, at least in part, by suppressing transcriptional activity, thereby decreasing downstream toxic protein burden.

### 3.4. Transcriptomic profiling reveals metabolic and functional reconfiguration associated with *K. rhizophila* (CDMP12)-mediated neuroprotection

To elucidate the molecular mechanisms underlying the observed neuroprotection and behavioral rescue upon feeding on CDMP12, we interrogated the global transcriptomic landscape of UA44 worms via RNA-sequencing. Comparative analysis revealed that CDMP12 drives a profound shift in gene expression, identifying 4,247 differentially expressed genes (DEGs) (p ≤ 0.05, |log_2_ FC| ≥ 0.5) (**Supplementary File 1**). This extensive modulation comprised 1,777 downregulated and 2,470 upregulated transcripts in UA44 (CDMP12) as compared to UA44 (OP50) (**Figures 4A**). Hierarchical clustering of the top 50 DEGs demonstrated a sharp segregation between the treatment and control groups, delineating a distinct transcriptional signature associated with the bacterial diet (**Figure 4B**). Collectively, these data indicate that *K. rhizophila* (CDMP12) induces a systemic transcriptomic reconfiguration, providing a robust dataset to decouple the specific pathways driving neuroprotection.

**Figure 4.**
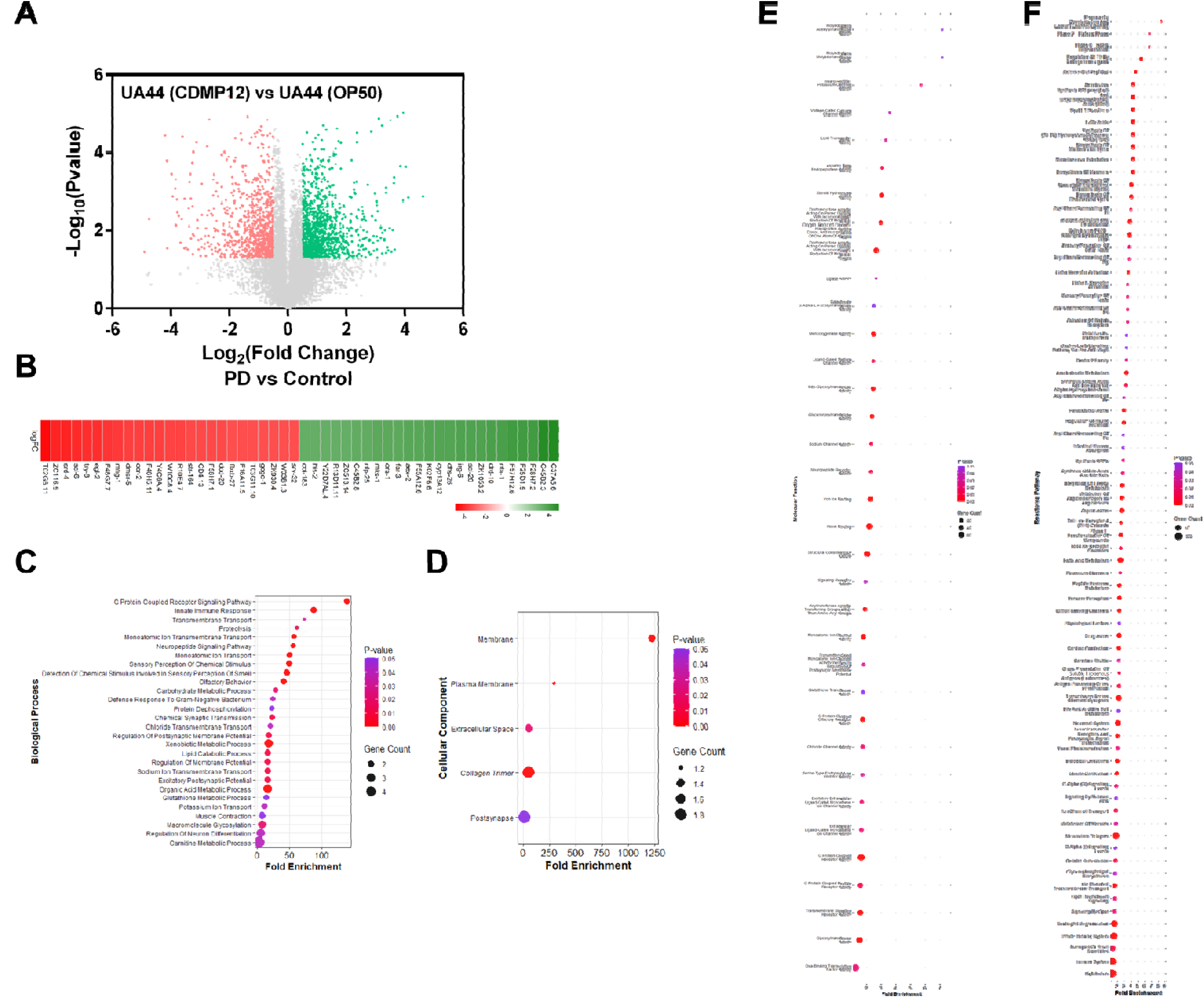
Transcriptomic reprogramming induced by *K. rhizophila* (CDMP12) in the UA44 strain. (A) Volcano plot depicting differentially expressed genes (DEGs) between UA44 (CDMP12) and UA44 (OP50). A total of 4,247 DEGs were identified (p ≤ 0.05, |log_2_ FC| ≥ 0.5), including 2,470 upregulated (green) and 1,777 downregulated (red) genes in UA44 (CDMP12). (B) Heatmap of the top 50 DEGs illustrating distinct transcriptional signatures between UA44 (CDMP12) and UA44 (OP50). Color scale represents log_2_ FC values. (C-F) Gene Ontology (GO) and Reactome pathway enrichment analysis of the DEGs using DAVID. Bubble plots showing the significant GO terms for (C) Biological Processes (D) Cellular Component, (E) Molecular Function, and (F) Reactome pathway enrichment.

To elucidate the biological relevance of the DEGs, comprehensive GO and pathway enrichment analyses were performed. GO analysis identified Carnitine Metabolic Process, Xenobiotic Metabolic Process, and Organic Acid Metabolic Process as the top three significantly enriched biological processes (**Figure 4C** **and Supplementary File 2**). Enrichment in cellular component terms primarily mapped to Collagen Trimer, Postsynapse membrane, and extracellular space (**Figure 4D** **and Supplementary File 2**). The top three significantly enriched molecular function categories included Molybdopterin Molybdotransferase Activity, Molybdopterin Adenylyltransferase Activity, and Inward Rectifier Potassium Channel Activity (**Figure 4E** **and Supplementary File 2**). Reactome pathway enrichment further revealed significant modulation of pathways associated with Presynaptic Depolarization and Calcium Channel Opening, Phase 0 - Rapid Depolarisation, and Phase 2 - Plateau Phase (**Figure 4F** **and Supplementary File 2**). These enriched categories highlight the crucial role of *K. rhizophila* (CDMP12) in modulating cellular metabolism, ion channel activity, and electrophysiological signaling pathways to provide neuroprotection in *C. elegans* PD model, UA44.

### 3.5. *K. rhizophila* (CDMP12) suppresses ferroptosis-mediated toxicity in the UA44 strain

An interesting finding from the Reactome pathway enrichment analysis was that many enriched pathways are associated with ferroptosis. Specifically, pathways including: Fatty Acid Metabolism & Fatty Acids, Arachidonate Metabolism, Slc-Mediated Transmembrane Transport, Metabolism of Lipids, Biological Oxidations, and Glycerophospholipid Biosynthesis & Acyl Chain Remodelling are known to modulate ferroptosis by regulating lipid substrate availability, membrane composition, and cellular redox homeostasis ^48–53^. Since ferroptosis is increasingly implicated in PD-associated neurodegeneration, these Reactome pathway enrichment results suggest that modulation of ferroptosis-related processes may contribute to the neuroprotective effects of *K. rhizophila* (CDMP12) against DA neuron degeneration in the UA44 model.

This hypothesis was supported by transcriptomic alterations in key ferroptosis-associated genes. Specifically, *smf-3*, the ortholog of human DMT1 (Divalent Metal Transporter 1), a principal mediator of cellular ferrous iron uptake is significantly downregulated in UA44 (CDMP12) (log_2_ FC= -1.76) (**Figure 5A**). Additionally, *ftn-1*, an ortholog of human FTH1 (ferritin heavy chain 1), was downregulated in UA44 (CDMP12) (log_2_ FC= -1.38) (**Figure 5A**). *Acs-4*, the ortholog of human ACSL4 (acyl-CoA synthetase long-chain family member 4) involved in fatty acid metabolism, was also significantly downregulated in UA44 (CDMP12) (log_2_ FC= -0.02), while *gpx-1*, the ortholog of human GPX4 (glutathione peroxidase 4), was upregulated (log_2_ FC= 0.4), reflecting a reinforced cellular defense against ferroptosis-associated lipid peroxidation (**Figure 5A**). Collectively, these coordinated transcriptional adaptations indicate that *K. rhizophila* (CDMP12) may confer neuroprotection by systematically suppressing ferroptosis pathways and fortifying cellular resistance to oxidative damage.

**Figure 5.**
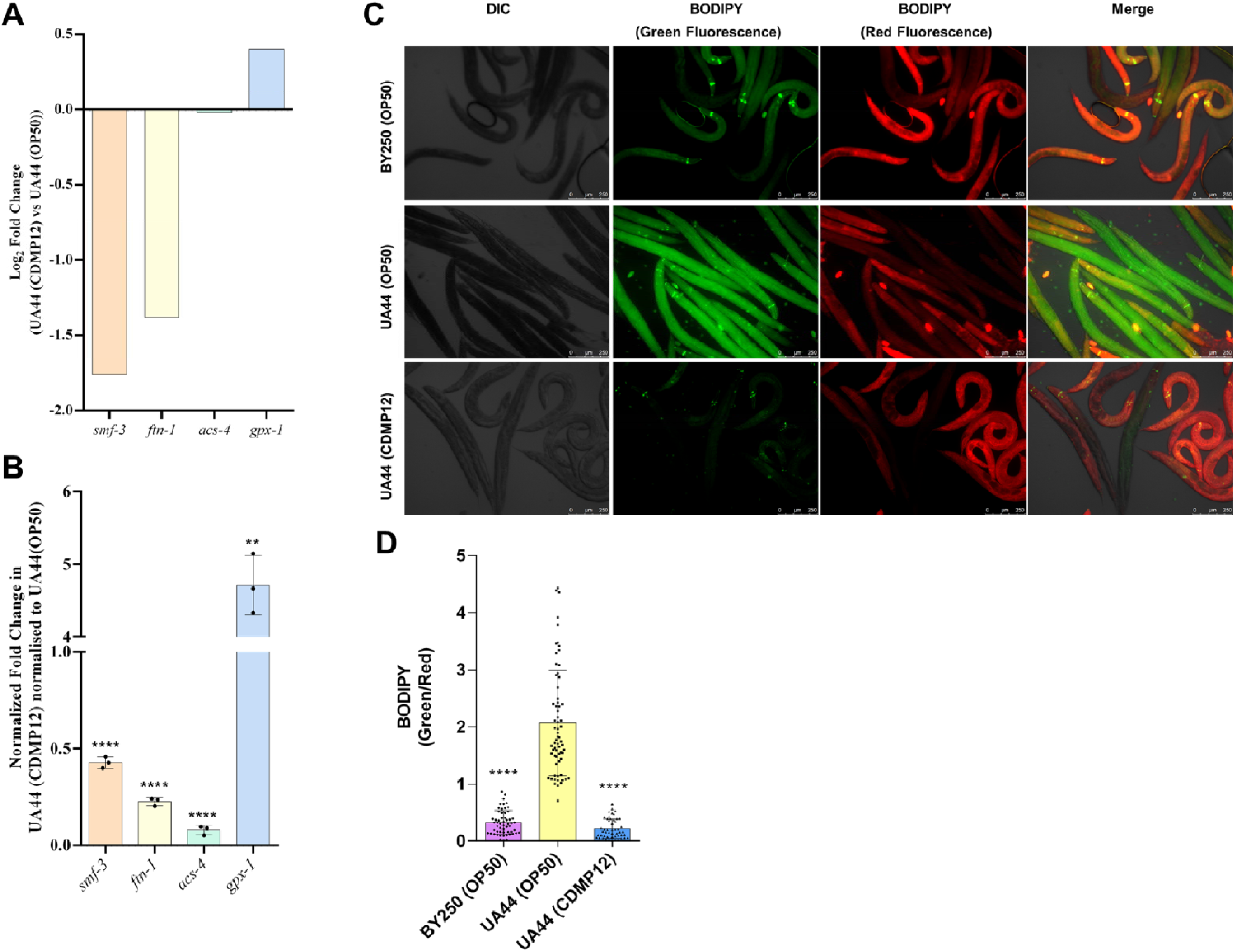
*K. rhizophila* (CDMP12) attenuates ferroptosis-related markers in the UA44 PD model. (A) Transcriptomic analysis revealed significant downregulation of *smf-3, ftn-1,* and *acs-4,* and upregulation of *gpx-1* in UA44 (CDMP12) worms compared to UA44 (OP50). (B) qRT-PCR validation confirmed reduced *smf-3, ftn-1,* and *acs-4* transcript levels and increased *gpx-1* transcript levels in UA44 (CDMP12) relative to UA44 (OP50), normalized to beta-actin (n=3 biological repeats; >500 worms per condition per repeat). (C) Representative BODIPY staining images showing oxidised lipids (green fluorescence), non-oxidised lipids (red fluorescence), and merged images. UA44 (OP50) worms displayed pronounced lipid peroxidation, which was markedly reduced in UA44 (CDMP12). Magnification=10X. (D) Quantification of BODIPY green/red fluorescence ratio confirmed reduced lipid peroxidation in UA44 (CDMP12) compared to UA44 (OP50) (n ≥ 30 worms per condition; pooled from at least three independent biological replicates). Data are presented as mean ± SD; **p < 0.01, ****p < 0.0001.

Using qRT-PCR, we validated the altered expression of these genes in UA44 (CDMP12) compared to the UA44 (OP50). Consistent with the transcriptomic data, *smf-3* (0.4 ± 0.02; p < 0.0001)*, ftn-1* (0.2 ± 0.01; p < 0.0001), and *acs-4* (0.08 ± 0.01; p < 0.0001) were downregulated, whereas *gpx-1* (5 ± 0.2; p < 0.01) was upregulated in UA44 (CDMP12) worms (**Figure 5B**).

To functionally validate whether these transcriptional changes translate to reduced lipid damage, a hallmark of ferroptosis, we next performed BODIPY 581/591 staining analysis on day three adult worms ^25,54^. The analysis demonstrated that UA44 (OP50) worms had a significant increase in the oxidized-to-non-oxidized lipid ratio (2.07 ± 0.11) when compared with BY250 (OP50) worms (0.32 ± 0.03; p < 0.0001) (**Figure 5C and D**). Crucially, UA44 (CDMP12) worms showed significant suppression in this ratio (0.22 ± 0.02; p < 0.0001) as compared to UA44 (OP50) (**Figure 5C and D**). These findings demonstrate that *K. rhizophila* (CDMP12) counters ferroptosis-driven oxidative stress, reducing a central pathogenic driver of PD-associated neurodegeneration.

## 4. Discussion

Originally discovered in plant roots, *Kocuria rhizophila* has since turned up in surprisingly diverse environments. While researchers have long valued it for stress tolerance, plant growth promotion, and producing antimicrobial compounds like bacteriocins, its therapeutic role in neurodegeneration has remained entirely unexplored ^26–31^. By demonstrating that strain CDMP12 protects against Parkinson’s-related neurodegeneration, this study expands the scope of marine bioprospecting into the domain of neuroprotection. *K. rhizophila* (CDMP12) exemplifies the potential of environmental microorganisms as viable reservoirs of bioactive molecules for addressing complex neurological disorders.

The preservation of DA neurons and the concurrent rescue of dopamine-dependent behaviors in CDMP12-fed UA44 worms provide evidence that microbial interventions can actively counteract neurotoxicity. In the broader landscape of PD research, these findings reinforce the hypothesis that dietary or commensal bacteria can exert systemic influence over neural circuitry. Rather than merely acting as a passive nutritional source, CDMP12 actively remodels host physiology. The restoration of functional neuro-sensory and motor behaviors indicates that the structural preservation is functionally meaningful, successfully re-establishing synaptic communication and neurotransmitter dynamics that are typically dismantled by alpha-synuclein toxicity.

A critical insight from our data is that CDMP12 reduces alpha-synuclein burden by suppressing its transcriptional activity at the mRNA level. In PD pathogenesis, intracellular alpha-synuclein accumulation creates a toxic gain-of-function that triggers neuroinflammation and mitochondrial dysfunction ^55^. By dampening transcription, CDMP12 cuts off the pathological cascade at its source. This suggests that bacterial metabolites or signaling molecules derived from *K. rhizophila* may interact with host regulatory pathways to suppress the expression of disease-associated transgenes.

Beyond inducing broad metabolic and functional reconfiguration, the transcriptomic alterations observed in this study highlight ferroptosis-associated processes as a potentially important component of CDMP12-mediated neuroprotection. Ferroptosis, an iron-driven form of regulated cell death, has emerged as a critical driver of PD progression ^56,57^. This pathological link is strongly supported by clinical evidence showing abnormal iron accumulation in the SNpc, which directly correlates with DA neuron loss and motor decline ^56,57^. Consequently, iron chelation has gained traction as a promising therapeutic strategy. For instance, the blood-brain barrier-permeable chelator deferiprone has demonstrated encouraging results in phase II clinical trials, successfully lowering nigral iron levels and alleviating motor symptoms in early-stage patients ^58,59^. Beyond synthetic chelators, natural polyphenols offer similar neuroprotection by sequestering excess iron, mitigating oxidative stress, and dampening alpha-synuclein aggregation ^60^. Intriguingly, the core molecular targets modulated by *K. rhizophila* (CDMP12) in this study align closely with established therapeutic strategies currently being explored to combat PD. For instance, therapeutic interventions targeting DMT1 and lipid peroxidation defense pathways have long been recognized as promising avenues for mitigating iron-induced neurotoxicity ^61^. Preclinical and clinical investigations into iron chelators together with pharmacological modulation of ACSL4 and enhancement of GPX4 activity, have consistently demonstrated robust protective effects against DA cell death ^62^. Building upon these established frameworks, our complementary evidence from transcriptomic profiling, qRT-PCR, and BODIPY lipid peroxidation assays reveals coordinated modulation of multiple ferroptosis-associated processes following *K. rhizophila* (CDMP12) treatment, suggesting that attenuation of ferroptotic stress may contribute to its neuroprotective effects.

*Smf-3* (*h*DMT1) encodes a divalent metal transporter involved in cellular ferrous iron uptake. In PD, increased DMT1 expression and activity have been associated with iron accumulation and oxidative stress, potentially contributing to an expanded labile iron pool and Fenton chemistry-mediated oxidative damage ^63^. In our study, CDMP12 significantly reduced *smf-3* mRNA expression, suggesting a potential shift toward reduced iron uptake. Similarly, *ftn-1*, the *C. elegans* ferritin heavy-chain ortholog involved in iron storage and homeostasis, was significantly downregulated following CDMP12 treatment. Together, these transcriptional changes suggest that CDMP12 modulates iron-homeostasis pathways, which may contribute to the observed reduction in ferroptosis-associated oxidative stress ^64,65^.

Beyond the observed changes in iron-homeostasis-associated genes, CDMP12 significantly reduced the expression of *acs-4*, the *C. elegans* ortholog of human ACSL4. ACSL4 is an important regulator of ferroptosis susceptibility, promoting the incorporation of polyunsaturated fatty acids into membrane phospholipids vulnerable to lipid peroxidation ^66^. Thus, reduced *acs-4* expression may decrease the availability of peroxidation-prone lipid substrates.

In parallel, CDMP12 significantly increased the expression of *gpx-1*, a *C. elegans* glutathione peroxidase associated with protection against lipid peroxidation, suggesting an enhanced transcriptional antioxidant response ^67^. Together, the downregulation of *acs-4* and upregulation of *gpx-1*, coupled with the marked reduction in BODIPY-detected lipid peroxidation, support an overall shift toward reduced lipid peroxidative stress following CDMP12 treatment.

Given the established interplay between alpha-synuclein accumulation, iron dyshomeostasis, oxidative stress, and lipid peroxidation in PD, the concurrent reduction in alpha-synuclein burden and ferroptosis-associated signatures following CDMP12 treatment raises the possibility that these processes are interconnected ^60^. However, further studies are required to establish their mechanistic relationship and determine whether modulation of alpha-synuclein directly influences ferroptotic vulnerability in this model.

Together, these findings establish *K. rhizophila* as a previously unrecognized marine-associated bacterium with neuroprotective efficacy in *C. elegans* models of PD, highlighting an exciting and largely untapped microbial resource for neurotherapeutics. The marine biome is increasingly recognized as a treasure trove of neuroactive compounds; for instance, brown-algal sulfated polysaccharides like fucoidan exhibit potent antioxidant and anti-inflammatory properties that safeguard DA neurons, while bromophenols derived from red algae act as multi-target agents capable of inhibiting monoamine oxidase and stimulating dopamine receptors ^68,69^. Within this expanding landscape of marine bioprospecting, *K. rhizophila* (CDMP12) introduces a fresh paradigm: a bacterial agent that simultaneously reigns in alpha-synuclein pathology, rescues DA neuronal survival, and suppresses ferroptosis-driven oxidative stress. This unique multi-pronged mechanism positions *K. rhizophila* (CDMP12) as a compelling and unconventional candidate for future translational neurodegenerative research.

Nevertheless, several limitations warrant note: *C. elegans* lacks the complex circuitry of mammalian brains, and while our data demonstrate a clear attenuation of iron-dependent lipid peroxidation and oxidative stress, they do not conclusively establish the direct inhibition of ferroptotic cell death without pharmacological confirmation using a canonical inhibitor like ferrostatin-1. Furthermore, given its marine origin and lack of established gut compatibility, we do not envision administering *K. rhizophila* as a live human probiotic; rather, it serves as a valuable biological source for discovering neuroactive postbiotic factors. Future investigations will focus on isolating these bacterial-derived metabolites, validating their efficacy in mammalian models, and incorporating genetic or pharmacological modulators to definitively map the ferroptotic contribution. Ultimately, by addressing core pathogenic drivers, iron dysregulation, lipid oxidation, and proteinopathy, this work demonstrates how environmental microbes can serve as innovative resources for developing disease-modifying therapies in PD.

## Abbreviations

PD: Parkinson’s disease
DA: Dopaminergic
SNpc: Substantia nigra pars compacta
NGM: Nematode Growth Medium
L1: First larval stage
DEGs: Differentially Expressed Genes
DAVID: Database for Annotation, Visualization, and Integrated Discovery
GO: Gene Ontology
FC: Fold change

## Data Availability Statement

The datasets generated for UA44 (Control) and UA44 (CDMP12) are available in the NCBI Gene Expression Omnibus (GEO) repository under accession code GSE312231 and GSE342134, respectively. The Reviewer Token is kdkloasqxrkxhab for accessing the GEO dataset GSE342134.

## CRediT Author Contribution Statement

**SS-** Investigation, Data curation, Validation, Formal analysis, Visualization, Conceptualization, Methodology, Writing – original draft.

**AD-** Investigation; Data curation; Formal analysis; Visualization, Writing - original draft.

**PY-** Investigation; Data curation

**MP-** Resources; Investigation; Methodology; Writing - original draft, Writing - review & editing.

**NK-** Data curation; Formal analysis; Writing - original draft; and Writing - review & editing.

**SV-** Conceptualization; Investigation; Data curation; Formal analysis; Methodology; Project administration; Resources; Supervision; Validation; Visualization; Writing - original draft; and Writing - review & editing.

## Generative AI statement

While preparing this work, the author(s) used Grammarly to improve language and readability. The author(s) reviewed and edited the content as needed and take(s) full responsibility for the content of the publication.

## Acknowledgements

SS is supported by the University Grants Commission, India. AD is supported by the Council of Scientific & Industrial Research, India. We gratefully acknowledge the Director of CSIR-CDRI for supporting this work by providing the essential research facilities.

## Ethics Statement

This study did not require approval from an institutional ethics committee, as it did not involve human participants, animal subjects, or sensitive data.

